# On the Calculation of Betweenness Centrality in Marine Connectivity Studies Using Transfer Probabilities

**DOI:** 10.1101/219881

**Authors:** Andrea Costa, Anne A. Petrenko, Katell Guizien, Andrea M. Doglioli

## Abstract

Betweenness has been used in a number of marine studies to identify portions of sea that sustain the connectivity of whole marine networks. Herein we highlight the need of methodological exactness in the calculation of betweenness when graph theory is applied to marine connectivity studies based on transfer probabilities. We show the inconsistency in calculating betweeness directly from transfer probabilities and propose a new metric for the node-to-node distance that solves it. Our argumentation is illustrated by both simple theoretical examples and the analysis of a literature data set.

## Introduction

In the last decade, graph theory has increasingly been used in ecology and conservation studies (Moilanen, 2011) and particularly in marine connectivity studies (e.g., Treml et al. 2008; Kininmonth et al. 2010a; Kininmonth et al. 2010b; Andrello et al. 2013; Rossi et al. 2014). Graphs are a mathematical representation of a network of entities (called nodes) linked by pairwise relationships (called edges). Graph theory is a set of mathematical results that permit to calculate different measures to identify nodes, or set of nodes, that play specific roles in a graph (e.g., Bondy and Murty 1976). Graph theory application to the study of marine connectivity typically consists in the representation of portions of sea as nodes. Then, the edges between these nodes represent transfer probabilities between these portions of sea.

Transfer probabilities estimate the physical dispersion of propagula (Andrello et al. 2013; Berline et al. 2014); (Jacobi et al. 2012; Jonsson et al. 2015), nutrients or pollutants (Doglioli et al., 2004), particulate matter (Mansui et al., 2015), or other particles either passive or interacting with the environment (see Ghezzo et al. 2015; Bacher et al. 2016; and references therein). As a result, graph theory already proved valuable in the identification of hydrodynamical provinces Rossi et al. (2014), genetic stepping stones (Rozenfeld et al., 2008), genetic communities (Kininmonth et al., 2010b), sub-populations (Jacobi et al., 2012), and in assessing Marine Protected Areas connectivity (Andrello et al., 2013).

In many marine connectivity studies, it is of interest to identify specific portions of sea where a relevant amount of the transfer across a graph passes through. A well-known graph theory measure is frequently used for this purpose: betweenness centrality. In the literature, high values of this measure are commonly assumed to identify nodes sustaining the connectivity of the whole network. For this reason a high value of betweenness has been used in the framework of marine connectivity to identify migration stepping stones (Treml et al., 2008), genetic gateways (Rozenfeld et al., 2008), and marine protected areas ensuring a good connectivity between them (Andrello et al., 2013).

Our scope in the present letter is to highlight some errors that can occur in implementing graph theory analysis. Especially we focus on the definition of edges when one is interested in calculating the betweenness centrality and other related measures. We also point out two papers in the literature in which this methodological inconsistency can be found: Kininmonth et al. (2010a) and Andrello et al. (2013).

In Materials and Methods we introduce the essential graph theory concepts for our scope. In Results we present our argument on the base of the analysis of a literature data set. In the last Section we draw our conclusions.

## Materials and Methods

A simple graph *G* is a couple of sets (*V, E*), where *V* is the set of nodes and *E* is the set of edges. The set *V* represents the collection of objects under study that are pair-wise linked by an edge *a_ij_*, with (*i, j*) *2 V*, representing a relation of interest between two of these objects. If *a_ij_* = *a_ji_*, ∀(*i, j*) *2 V*, the graph is said to be ‘undirected’, otherwise it is ‘directed’. The second case is the one we deal with when studying marine connectivity, where the edges’ weights represent the transfer probabilities between two zones of sea (e.g., Kininmonth et al. 2010a; Kininmonth et al. 2010b; Andrello et al. 2013; Rossi et al. 2014).

If more than one edge in each direction between two nodes is allowed, the graph is called multi-graph. The number of edges between each pair of nodes (*i, j*) is then called multiplicity of the edge linking *i* and *j*.

The in-degree of a node *k*, *deg*^+^(*k*), is the sum of all the edges that arrive in *k*: *deg*^+^(*k*) = *Σ_i_ a_ik_*. The out-degree of a node *k*, *deg^−^*(*k*), is the sum of all the edges that start from *k*: *deg^+^*(*k*) = *Σ_j_ a_kj_*. The total degree of a node *k*, *deg*(*k*), is the sum of the in-degree and out-degree of *k*: *deg*(*k*) = *deg*^+^(*k*) + *deg^−^*(*k*).

In a graph, there can be multiple ways (called paths) to go from a node *i* to a node *j* passing by other nodes. The weight of a path is the sum of the weights of the edges composing the path itself. In general, it is of interest to know the shortest or fastest path *σ_ij_* between two nodes, i.e. the one with the lowest weight. But it is even more instructive to know which nodes participate to the greater numbers of shortest paths. In fact, this permits to measure the influence of a given node over the spread of information through a network. This measure is called betweenness value of a node in the graph. The betweenness value of a node *k*, *BC*(*k*), is defined as the fraction of shortest paths existing in the graph, *σ_ij_*, with *i* ≠ *j*, that effectively pass through *k*, *σ_ij_*(*k*), with *i* ≠ *j* ≠ *k*:

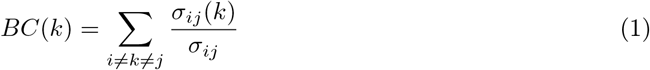

with (*i, j, k*) *∈ V*. Note that the subscript *i* ≠ *k* ≠ *j* means that betweenness is not influenced by direct connections between the nodes. Betweenness is then normalized by the total number of possible connections in the graph once excluded node *k*: (*N −* 1)(*N −* 2), where *N* is the number of nodes in the graph, so that 0 ≤ *BC* ≤ 1.

Although betweenness interpretation is seemingly straightforward, one must be careful in its calculation. In fact betweenness interpretation is sensitive to the node-to-node metric one chooses to use as edge weight. If, as frequently the case of the marine connectivity studies, one uses transfer probabilities as edge weight, betweenness loses its original meaning. Based on additional details –personally given by the authors of Kininmonth et al. (2010a) and Andrello et al. (2013)– on their methods, this was the case in those studies. In those cases, edge weight would decrease when probability decreases and the shortest paths would be the sum of edges with lowest value of transfer probability. As a consequence, high betweenness would be associated to the nodes through which a high number of improbable paths pass through. Exactly the opposite of betweenness original purpose. Hence, defining betweenness using Equation 1 (the case of Kininmonth et al. 2010a and Andrello et al. 2013) leads to an inconsistency that affects the interpretation of betweenness values. Alternative definitions of betweenness accounting for all the paths between two nodes and not just the most probable one have been proposed to analyze graphs in which the edge weight is a probability Newman (2005) and avoid the above inconsistency.

Herein, we propose to solve the inconsistency when using the original betweenness definition of transfer probabilities by using a new metric for the edge weights instead of modifying the betweenness definition. The new metric transforms transfer probabilities *a_ij_* into a distance in order to conserve the original meaning of betweenness, by ensuring that a larger transfer probability between two nodes corresponds to a smaller node-to-node distance. Hence, the shortest path between two nodes effectively is the most probable one. Therefore, high betweenness is associated to the nodes through which a high number of probable paths pass through.

In the first place, in defining the new metric, we need to reverse the order of the probabilities in order to have higher values of the old metric *a_ij_* correspond to lower values of the new one. In the second place we also consider three other facts: (i) transfer probabilities *a_ij_* are commonly calculated with regards to the position of the particles only at the beginning and at the end of the advection period; (ii) the probability to go from *i* to *j* does not depend on the node the particle is coming from before arriving in *i*; and (iii) the calculation of the shortest paths implies the summation of a variable number of transfer probability values. Note that, as the *a_ij_* values are typically calculated on the base of the particles’ positions at the beginning and at the end of a spawning period, we are dealing with paths whose values are calculated taking into account different numbers of generations. Therefore, the transfer probabilities between sites are independent from each other and should be multiplied by each other when calculating the value of a path. Nevertheless, the classical algorithms commonly used in graph theory analysis calculate the shortest paths as the summation of the edges composing them (e.g., the Dijkstra algorithm, Djikstra 1959; or the Brandes algorithm, Brandes 2006). Therefore, these algorithms, if directly applied to the probabilities at play here, are incompatible with their independence.

A possible workaround could be to not use the algorithms in Djikstra (1959) and Brandes (2006) and use instead the 10*^th^* algorithm proposed in Brandes (2008). Therein, the author suggests to define the betweenness of a simple graph via its interpretation as a multigraph. He then shows that the value of a path can be calculated as the product of the multiplicities of its edges. When the multiplicity of an edge is set equal to the weight of the corresponding edge in the simple graph, one can calculate the value of a path as the product of its edges’ weights *a_ij_*. However, this algorithm selects the shortest path on the basis of the number of steps (or hop count) between a pair of nodes (Breadth-First Search algorithm, Moore 1959). This causes the algorithm to fail in identifying the shortest path in some cases. For example, in Figure 1 it would identify the path ACB (2 steps with total probability 1 *×* 10*^−^*8) when, instead, the most probable path is ADEB (3 steps with total probability 1 *×* 10*^−^*6). See Table 1 for more details.

**Figure 1:**
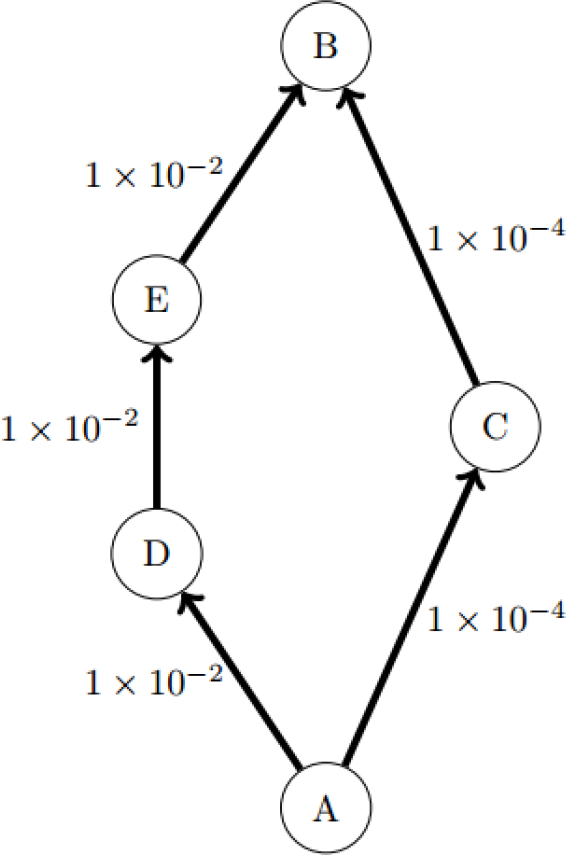
Example of graph in which the 10*^th^* algorithm in Brandes (2008) would fail to identify the shortest path between A and B (ADEB) when using *a_ij_* as metric.

**Table 1:**
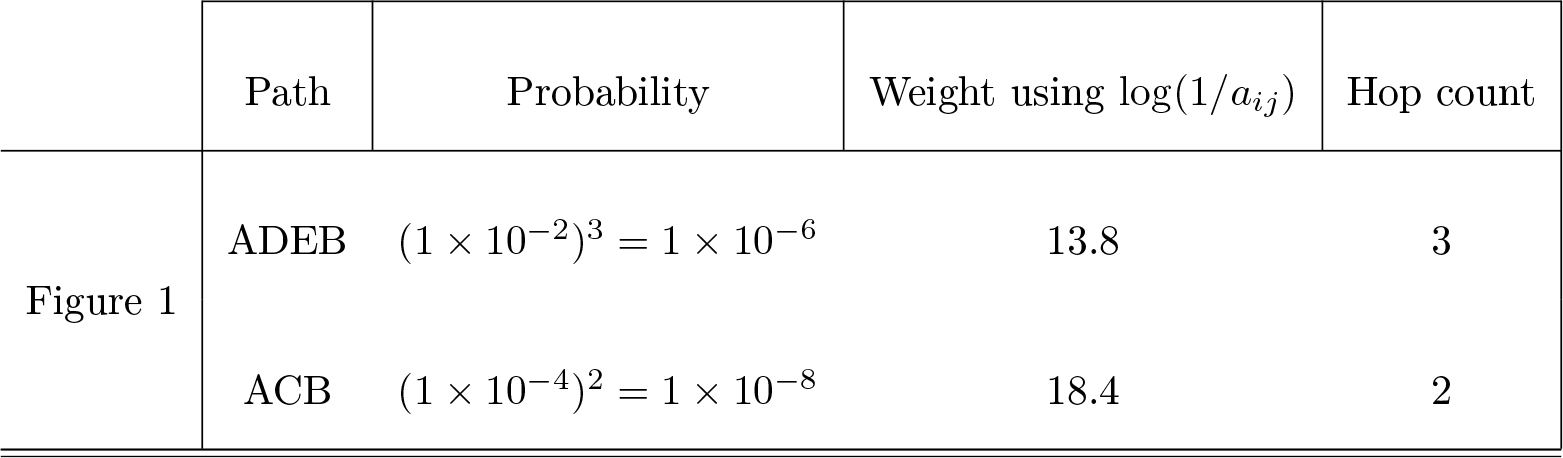
Paths and respective probabilities, weights and hop count for the graph in Figure 1.

However, by changing the metric used in the algorithms, it is possible to calculate the shortest path in a meaningful way with the algorithms in Djikstra (1959) and Brandes (2006). In particular, we propose to define the weight of an edge between two nodes *i* and *j* as:

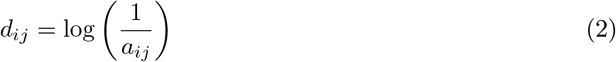

This definition is the composition of two functions: *h*(*x*) = 1*/x* and *f*(*x*) = log(*x*). The use of *h*(*x*) allows one to reverse the ordering of the metric in order to make the most probable path the shortest. The use of *f*(*x*), thanks to the basic properties of logarithms, allows the use of classical shortest-path finding algorithms while dealing correctly with the independence of the connectivity values. In fact, we are *de facto* calculating the value of a path as the product of the values of its edges.

It is worth mentioning that the values *d_ij_* = *∞*, coming from the values *a_ij_* = 0, do not influence the calculation of betweenness values via the Dijkstra and Brandes algorithms. Note that *d_ij_* is additive: 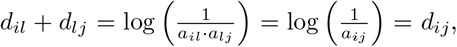 for any (*i, l, j*) ∞ *V* thus being suitable to be used in conjunction with the algorithms proposed by Djikstra (1959) and Brandes (2006). Also, note that both *a_ij_* and *d_ij_* are dimensionless. Equation 2 is the only metric that allows to consistently apply the Dijkstra and Brandes algorithms to transfer probabilities. Other metrics would permit to make the weight decrease when probability increases: for example, 1 *+ a_ij_*, 1*/a_ij_*, *−a_ij_*, log(1 *+ a_ij_*). However, the first three ones do not permit to account for the independence of the transfer probabilities along a path. Furthermore, log(1 *+ a_ij_*) takes negative values as 0 *a_ij_* 1. Therefore, it cannot be used to calculate shortest paths because the algorithms in Djikstra (1959) and Brandes (2006) would either endlessly go through a cycle (see Figure 2a and Table 2) or choose the path with more edges (see Figure 2b and Table 2), hence arbitrarily lowering the value of the paths between two nodes.

**Figure 2:**
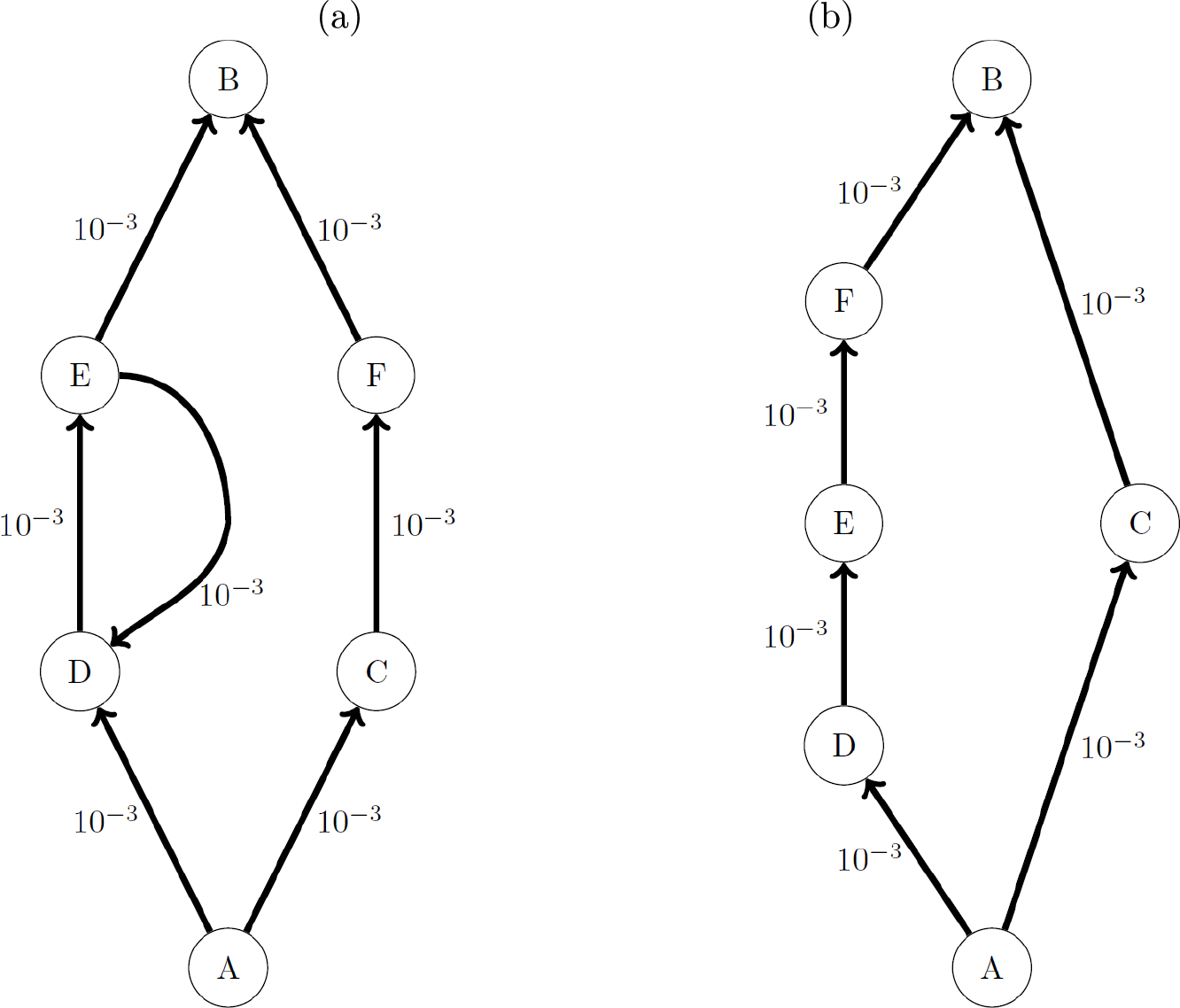
a) Example of network in which the metric log(1 − *a_ij_*) would fail because of a cycle (ED). b) Example of network in which the metric log(1 − *a_ij_*) would fail by taking the longest path possible (ADEFB instead of ACB).

**Table 2:**
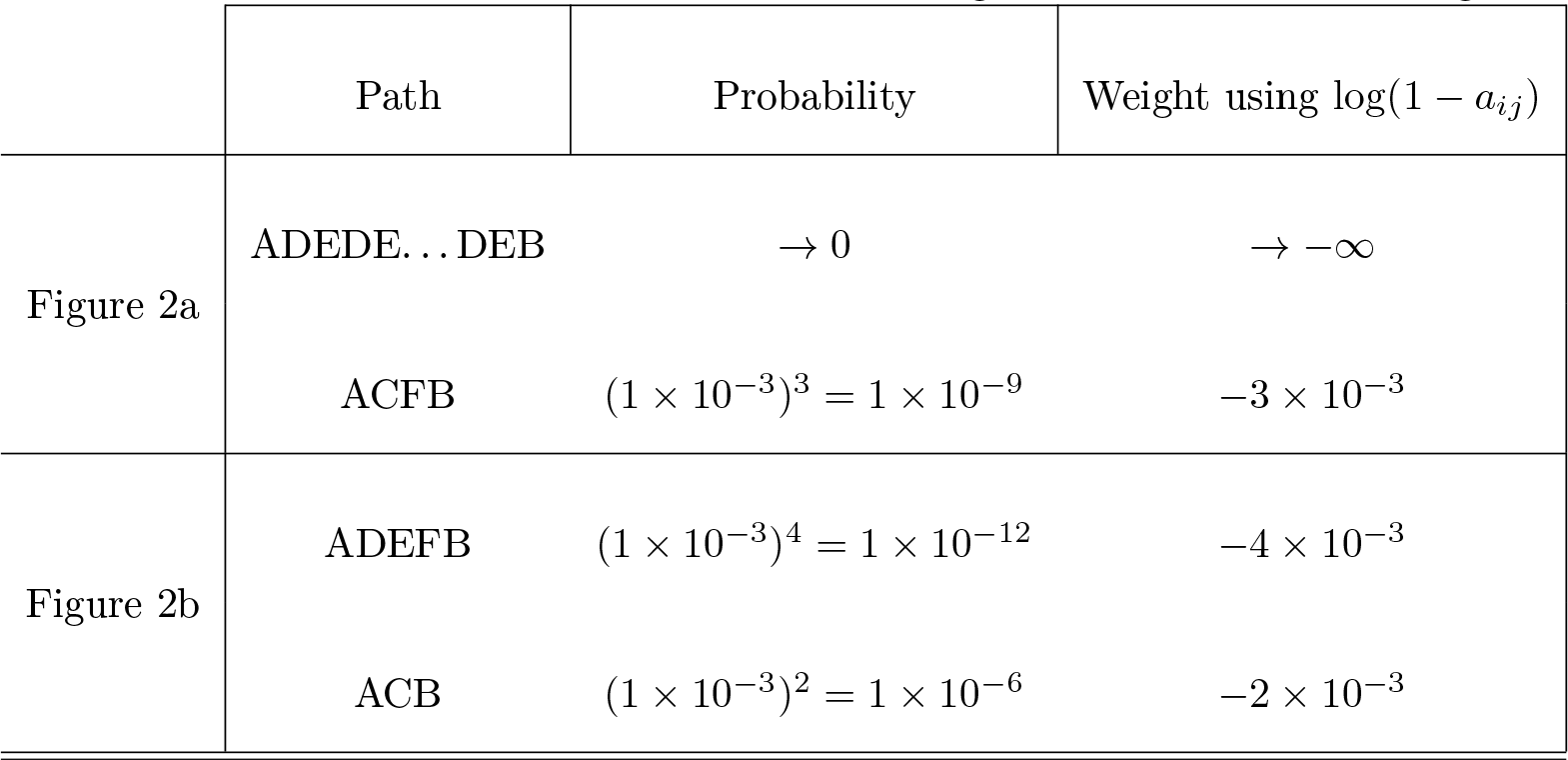
Paths and respective probabilities and weights for the networks in Figure 2.

## Results

The consequences of the use of the raw transfer probability (*a_ij_*) rather than the distance we propose (*d_ij_*) are potentially radical. To show this, we used 20 connectivity matrices calculated for Guizien et al. (2014). They were calculated from Lagrangian simulations using a 3D circulation model with a high horizontal resolution of 750 *m* (Marsaleix et al., 2006). Spawning was simulated by releasing 30 particles in the center of each of 32 reproductive sites (hereafter identified as nodes) for benthic polychaetes alongshore the Gulf of Lion (NW Mediterranean Sea), on the 30 *m* isobath, every hour from January 5 until April 13 in 2004 and 2006. Note that the connectivity matrices’ values strongly depend on the circulation present in the Gulf during the period of the dispersal simulations. The typical circulation of the Gulf of Lion is a westward current regime Millot (1990). This was the case of matrices #7,#11, #12, #15, #17. However, other types of circulation are often observed. In particular matrix #1 was obtained after a period of reversed (eastward) circulation. Indeed, this case of circulation is less frequent than the westward circulation (Petrenko et al., 2008). Matrices #14, #10 and #13 correspond to a circulation pattern with an enhanced recirculation in the center of the gulf. Finally, matrices #2, #3, #5, #6, #8, #9, #14, #16, #18, #19, #20 correspond to a rather mixed circulation with no clear pattern. The proportions of particles coming from an origin node and arriving at a settlement node after 3, 4 and 5 weeks were weight-averaged to compute a connectivity matrix for larvae with a competency period extending from 3 to 5 weeks.

As an example, in Fig 3 we show the representation of the graph corresponding to matrix #7. The arrows starting from a node *i* and ending in a node *j* represent the direction of the element *a_ij_* (in Fig 3a) or *d_ij_* (in Fig 3b). The arrows’ color code represents the magnitude of the edges’ weights. The nodes’ color code indicates the betweenness values calculated using the metric *a_ij_* (in Fig 3a) or *d_ij_* (in Fig 3b).

**Figure 3:**
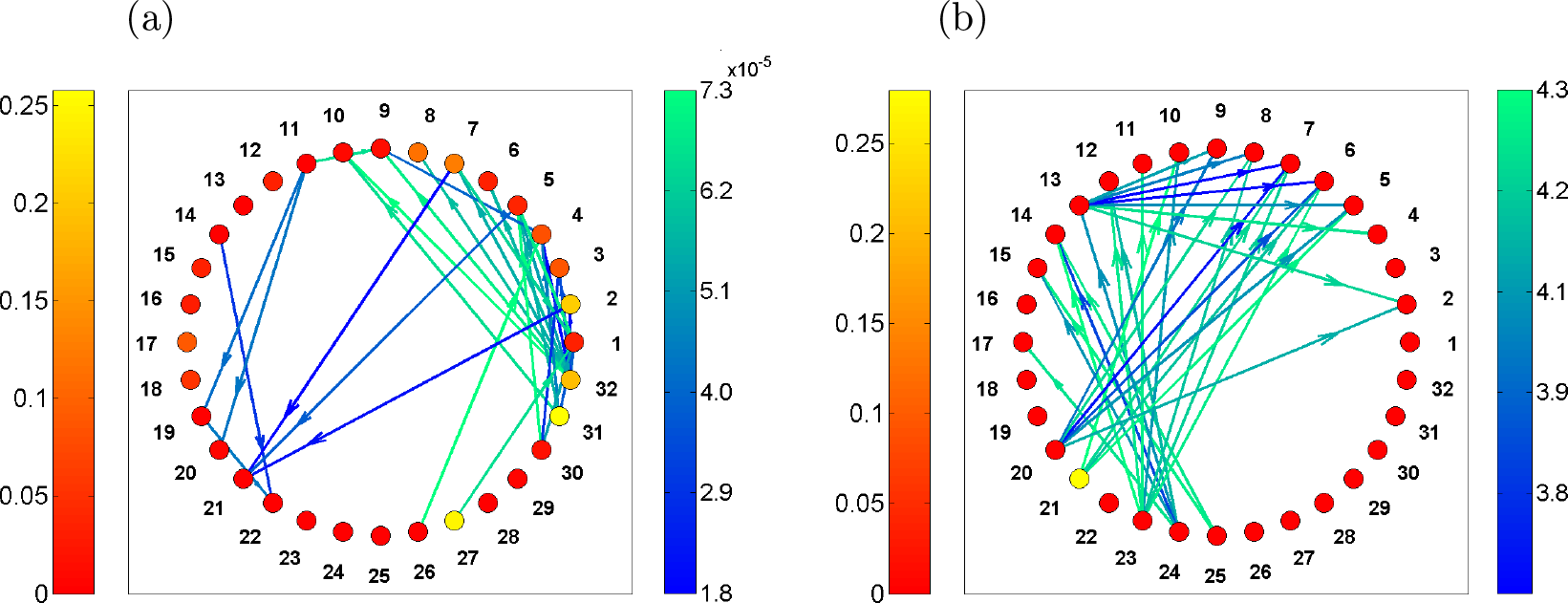
Representation of matrix matrix #7 from Guizien et al. (2014), the right side colorbars indicate the metric values. a) Results obtained by using *a_ij_* as edge weight, b) results obtained by using *d_ij_* as edge weight. In a) the lowest 5% of edges weights are represented. In b) the lowest 5% of edges weights are represented. Note the change in the colorbars’ ranges.

In Fig 3a the edges corresponding to the lower 5% of the weights *a_ij_* are represented. These are the larval transfers that, though improbable, are the most influential in determining high betweenness values when using *a_ij_* as metric. In Fig 3b the edges corresponding to the lower 5% of the weights *d_ij_* are represented. These are the most probable larval transfers that —correctly— are the most influential in determining high betweenness values when using *d_ij_* as metric. While in figure Fig 3a the nodes with highest betweenness are the nodes 31 (0.26), 27 (0.25) and 2 (0.21); in Fig 3b the nodes with highest betweenness are nodes 21 (0.33), 20 (0.03) and 29 (0.03).

Furthermore, it is expected to have a positive correlation between the degree of a node and its betweenness (e.g., Valente et al. 2008 and Lee 2006). However, we find that the betweenness values, calculated on the 20 connectivity matrices containing *a_ij_*, have an average correlation coefficient of *−*0.42 with the total degree, *−*0.42 with the in-degree, and *−*0.39 with the out-degree. Instead, betweenness calculated with the metric of Equation 2 has an average correlation coefficient of 0.48 with the total degree, 0.45 with the in-degree, and a not significant correlation with the out-degree (p-value > 0.05).

As we show in Fig 4, betweenness values of the 32 nodes calculated using the two node-to-node distances *a_ij_* and log(1*/a_ij_*) are drastically different between each other. Moreover, in 10 out of 20 connectivity matrices, the correlation between node ranking based on betweenness values with the two metrics were not significant. In the 10 cases it was (p-value < 0.05), the correlation coefficient was lower than 0.6 (data not shown). Such partial correlation is not unexpected as the betweenness of a node with a lot of connections could be similar when calculated with *a_ij_* or *d_ij_* if among these connections there are both very improbable and highly probable ones, like in node 21 in the present test case. Furthermore, it is noticeable that if one uses the *a_ij_* values (Fig 4a), the betweenness values are much more variable than the ones obtained using *d_ij_* (Fig 4b). This is because, in the first case, the results depend on the most improbable connections that, in the ocean, are likely to be numerous and unsteady.

**Figure 4:**
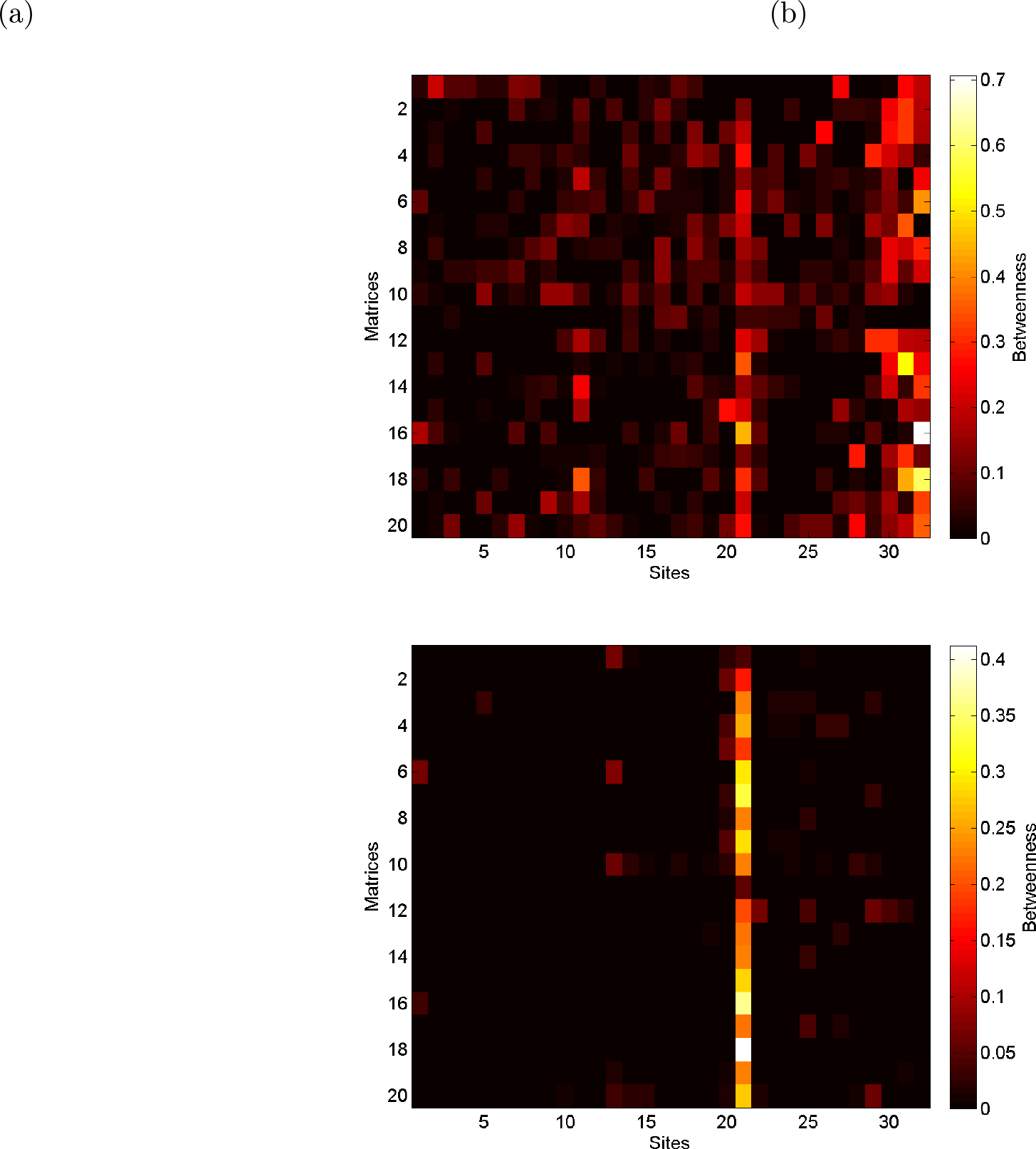
Betweenness values for the 32 sites in the Gulf of Lion using 20 different connectivity matrices obtained with Lagrangian simulations by Guizien et al. (2014). a) Results obtained by using *a_ij_* as edge weight; b) results obtained by using *d_ij_*. Note the change in the colorbars’ ranges.

## Conclusion

We highlighted the need of methodological exactness inconsistency in the betweenness calculation when graph theory to marine transfer probabilities. Indeed, the inconsistency comes from the need to reverse the probability when calculating shortest paths. If this is not done, one considers the most improbable paths as the most probable ones. We showed the drastic consequences of this methodological error on the analysis of a published data set of connectivity matrices for the Gulf of Lion (Guizien et al., 2014).

On the basis of our study, it may be possible that results in Kininmonth et al. (2010a) and Andrello et al. (2013) might also be affected. A re-analysis of Kininmonth et al. (2010a) would not affect the conclusions drawn by the authors about the small-world characteristics of the Great Barrier Reef as that is purely topological characteristics of a network. About Andrello et al. (2013), according to Marco Andrello (personal communication), due to the particular topology of the network at study, the change of the high-betweenness nodes does not drastically affect the main conclusions presented in Andrello et al. (2013), even if local changes of betweenness values are noticeable. To solve the highlighted inconsistency, we proposed the use of a node-to-node metric that provides a meaningful way to calculate shortest paths and —as a consequence— betweenness, when relying on transfer probabilities issued from Lagrangian simulations and the algorithm proposed in Djikstra (1959) and Brandes (2006). The new metric permits to reverse the probability and to calculate the value of a path as the product of its edges and to account for the independence of the transfer probabilities. Moreover, this metric is not limited to the calculation of betweenness alone but is also valid for the calculation of every graph theory measure related to the concept of shortest paths: for example, shortest cycles, closeness centrality, global and local efficiency, and average path length (Costa et al., In Prep.).

## Acknowledgments

The authors thank Dr. S.J. Kininmonth and Dr. M. Andrello for kindly providing the code they used for the betweenness calculation in their studies. The first author especially thanks Dr. R. Puzis for helpful conversations. Andrea Costa was financed by a MENRT Ph.D. grant. The research leading to these results has received funding from the European Union’s Seventh Framework Programme for research, technological development and demonstration under Grant Agreement No. 287844 for the project ‘Towards COast to COast NETworks of marine protected areas (from the shore to the high and deep sea), coupled with sea-based wind energy potential’ (COCONET).

## References

Andrello, M., Mouillot, D., Beuvier, J., Albouy, C., Thuiller, W., Manel, S., 2013. Low connectivity between Mediterranean Marine Protected Areas: a biophysical modeling approach for the dusky grouper: Epinephelus Marginatus. PLoS ONE 8, 1–15.

Bacher, C., Filgueira, R., Guyondet, T., 2016. Probabilistic approach of water residence time and connectivity using Markov chains with application to tidal embayments. J. Marine Syst 153, 25–41.

Berline, L., Rammou, A.M., Doglioli, A., Molcard, A., Petrenko, A., 2014. A connectivity-based ecoregionalization of the Mediterranean Sea. PLoS ONE 9.

Bondy, J., Murty, U., 1976. Graph theory with applications. Elsevier Science Publishing.

Brandes, U., 2006. A faster algorithm for betweenness centrality. J. Math. Sociol. 13, 163–177.

Brandes, U., 2008. On variants of shortest-path betweenness centrality and their generic computation. Soc. Networks 30, 1–22.

Costa, A., Doglioli, A., Guizien, K., Petrenko, A., In Prep. Tuning the interpretation of graph theory measures in analyzing marine larval connectivity.

Djikstra, E., 1959. A note on two problems in connection with graphs. Numerische Mathematik 1, 269–271.

Doglioli, A., Magaldi, M., ans S. Tucci, L.V., 2004. Development of a numerical model to study the dispersion of wastes coming from a marine fish farm in the Ligurian Sea (Western Mediterranean. Aquaculture 231, 215–235.

Ghezzo, M., Pascalis, F.D., Umgiesser, G., Zemlys, P., Sigovini, M., Marcos, C., Perez-Ruzafa, A., 2015. Connectivity in three European coastal lagoons. Estuar. Coasts 38, 1764–1781.

Guizien, K., Belharet, M., Moritz, C., Guarini, J., 2014. Vulnerability of marine benthic metapopulations: implications of spatially structured connectivity for conservation practice. Divers. Distrib. 20, 1392–1402.

Jacobi, M., André, C., Doos, K., Jonsson, P., 2012. Identification of subpopulations from connectivity matrices. Ecography 35, 31–44.

Jonsson, P., Jacobi, M., Moksnes, P.O., 2015. How to select networks of marine protected areas for multiple species with different dispersal strategies. Divers. Distrib. 22, 1–13.

Kininmonth, S., De’ath, M., Possingham, H., 2010a. Graph theoretic topology of the Great but small Barrier Reef world. Theor. Ecol. 3, 75–88.

Kininmonth, S., van Hopper, M., Possingham, H., 2010b. Determining the community structure of the coral *Seriatopora hystrix* from hydrodynamic and genetic networks. Ecol. Model. 221, 2870–2880.

Lee, C.Y., 2006. Correlations among centrality measures in complex networks. ArXiv Physics e-prints arXiv:physics/0605220.

Mansui, J., Molcard, A., Ourmieres, Y., 2015. Modelling the transport and accumulation of floating marine debris in the Mediterranean basin. Marine Poll. Bull. 91, 249–257.

Marsaleix, P., Auclair, F., Estournel, C., 2006. Considerations on open boundary conditions for regional and coastal ocean models. Ocean Technol. 23, 1604–1613.

Millot, C., 1990. The Gulf of Lion’s hydrodynamics. Cont. Shelf Res. 10, 885–894.

Moilanen, A., 2011. On the limitations of graph-theoretic connectivity in spatial ecology and conservation. Journal of Applied Ecology 48, 1543–1547.

Moore, E., 1959. The shortest path through a maze. Int. Symp. on Th. of Switching 1959, 285–292.

Newman, M., 2005. A measure of betweenness centrality based on random walks. Soc. Networks 27, 39–54.

Petrenko, A., Dufau, C., Estournel, C., 2008. Barotropic eastward currents in the western Gulf of Lion, north-western Mediterranean Sea, during stratified conditions. Journal of Marine Systems 74, 406–428.

Rossi, V., ans A.A. Lopez Cristobal, E.S.G., Hernandez-Garcia, E., 2014. Hydrodynamic provinces and oceanic connectivity from a transport network help designing marine reserves. Geoph. Res. Lett. 93, 2883–2891.

Rozenfeld, A., Arnaud-Haond, S., Hernandez-Garcia, E., Eguiluz, V., Serrao, E., Duarte, C., 2008. Network analysis identifies weak and strong links in a metapopulation system. Proc. Natl. Acad. Sci. USA 105, 18824–18829.

Treml, E., Halpin, P., Urban, D., Pratson, L., 2008. Modeling population connectivity by ocean currents, a graph theoretic approach for marine conservation. Lansc. Ecol. 23, 19–36.

Valente, T., Corognes, K., Lakon, C., Constenbader, E., 2008. How correlated are network centrality measures? Connections 28, 16–26.

